# Homologs of SD6 and ICE2 from rice may be involved in regulation of ABA in woody perennial buds

**DOI:** 10.1101/2023.10.25.563948

**Authors:** Zhaowan Shi, Sonika Pandey, Tamar Halaly-Basha, David W. Galbraith, Etti Or

**Author notes:** **Corresponding author**: Etti Or. Zhaowan Shi ^a^ Sonika Pandey ^a^ Tamar Halaly-Basha ^a^ David W. Galbraith ^b^ Etti Or ^a^.

## Abstract

The availability of ABA, a central component in the regulation of the dormancy cycle in grapevine buds, is controlled by coordinated and opposite regulation of NCED and ABA8OX expression, as shown during natural dormancy release and following treatment with Hydrogen Cyanamide (HC). This implies the existence of a shared regulatory entity, which serves as an upstream switch.

A molecular switch for integrated and opposite regulation of NCED and ABA8OX was recently described in rice, involving a pair of bHLH transcription factors (OsSD6-OsICE2) that directly regulate ABA8OX3 expression and indirectly regulate NCED2 expression, by direct regulation of the expression of the NCED repressor OsbHLH048.

Here, we tested whether expression of the *Vitis* homologs of the rice SD6 and ICE2 are regulated by dormancy release stimuli, and whether the direction of regulation agrees with that of *ABA8OX*.

Treatment with two independent stimuli of bud break (HC and hypoxia), as well as natural dormancy release, resulted in upregulation of OsSD6 homologs and down regulation of OsICE2 homologs, in agreement with the rice model. In unexpected contrast, the homolog of OsbHLH048 was down-regulated.

Our results suggest a grapevine model in which (1) the homologs of *OsSD6* and *OsICE2* act as direct activators and repressors of *ABA8OX3* expression, as for rice, (2) they have opposed effects on the expression of an OsbHLH048 homolog, which serves as direct activator of NCED expression, as for Arabidopsis, and (3) together they act as a switch that allows removal of ABA repression, followed by meristem reactivation and bud break.

**HIGHLIGHT:** A molecular switch for integrated and opposite regulation of NCED and ABA8OX in rice seeds, operated by three bHLH transcription factors, is conserved in grapevine buds and regulates dormancy release

## INTRODUCTION

Hydrogen cyanamide (HC) initiates a biochemical cascade that leads to effective dormancy release of grape buds, and its application is mandatory for commercial table grape production in warm-winter regions. Our former studies provided a working model of the molecular pathways and mechanisms that might regulate dormancy release of grape bud. In this model, perturbation of cytochrome-pathway activity in mitochondria leads to a transient respiratory and oxidative stress, resulting in an energy crisis that is accompanied by an oxidative burst. To address this crisis, glycolysis, anaerobic respiration, and cellular antioxidant machinery are coordinately induced. This reprogramming, which under aerobic environmental conditions mimics the hypoxia response, affects ethylene and abscisic acid (ABA) metabolism and/or signaling, leading to activation of catabolic pathways required to supply energy from alternative sources, removal of ABA repression and reactivation of shoot apical meristem growth (Ophir *et al*., 2009; Shi *et al*., 2020 and references within). Various predictions of this model were experimentally confirmed, as formerly reported (Ophir *et al*., 2009; Pérez *et al*., 2009; Shi *et al*., 2020; Shi *et al*., 2018; Vergara *et al*., 2012; Zheng *et al*., 2018a; Zheng *et al*., 2018b; Zheng *et al*., 2015).

Of special importance in the current context are the findings that are related to the potential role of ABA. Among those are the following outcomes: (1) across the natural bud dormancy cycle in grapes, endogenous ABA levels increase at the onset of dormancy, and decrease towards dormancy release (Koussa, 1994, Or *et al*., 2000); (2) exogenous ABA inhibits dormancy release of grapevine buds, and attenuates the enhancing effects of various artificial dormancy release stimuli, such as HC, sodium azide (AZ), and heat shock (HS) (Zheng *et al*., 2018b; Zheng *et al*., 2015); (3) HC is involved in diminishing the ABA repression potential via modification of ABA metabolism. It coordinately decreases the level of the *VvNCED1* transcript, and increases that of the *VvA8H-CYP707A4* transcript (Zheng *et al*., 2018b; Zheng *et al*., 2015); (4) the mode of regulation suggested by the changes recorded in the transcript levels is in agreement with the final outcome in metabolite levels, as reflected by the decrease in endogenous bud ABA and the parallel increase in ABA-degradation products in HC-treated buds, (5) similar coordination was evident across the natural dormancy cycle, with a significant decrease in the level of *VvNCED1* transcript being accompanied by a simultaneous sharp increase in the levels of *VvA8H-CYP707A4* at deepest dormancy status. The level of ABA catabolites was significantly increased concomitantly with the sharp increase in the *VvA8H-CYP707A4* transcript level. These findings suggest the existence of a defined developmental window in the natural bud dormancy cycle, within which ABA can play a regulatory role in dormancy maintenance; (6) the central role for ABA in maintaining grape bud dormancy was further validated by over-expression of *VvA8H-CYP707A4* gene, that led to decreased bud ABA levels and advances bud break (Zheng *et al*., 2018b).

Such a central role for ABA is in agreement with the effect of ABA on bud break in willow (Barros and Neill, 1989), apple (Dutcher and Powell, 1972), pear (Tamura *et al*., 2002), kiwi (Lionakis and Schwabe, 1984), sour cherry (Mielke and Dennis, 1978) and poplar (Tylewicz *et al*., 2018). It is also in agreement with (1) the parallel increase in *NCED* expression and ABA level during dormancy induction of poplar apical buds (Ruttink *et al*., 2007; Rohde *et al*., 2002) and (2) the positive correlation of upregulation of *StNCED2* and *StCYP707A1*, with increase and decrease of ABA content during progression of the natural dormancy cycle in potato tuber meristems, respectively (Destefano-Beltrán *et al*., 2006a; Destefano-Beltrán *et al*., 2006b). It is also in agreement with previous reports that presented the negative effects of exogenous ABA on seed germination (Rubio *et al*., 2009; Santiago *et al*., 2009; Ye *et al*., 2012), and the negative effect of ABA8’OH levels on seed dormancy. For example, Arabidopsis *cyp707a2* mutant presents increased levels of ABA in dry and imbibed seeds, and reduced germination (Okamoto *et al*., 2006). Similarly, transgenic *HvABA8’OH1* RNAi barley grains contain higher ABA levels and display increased dormancy (Gubler *et al*., 2008). Constitutive expression of *AtCYP707A1* gene in transgenic Arabidopsis plants resulted in a decreased ABA content in mature dry seeds, and a much shorter after-ripening period to overcome dormancy (Millar *et al*., 2006). This commonality suggests the existence of conserved primary regulatory blocks that control the activity of the meristem in dormant buds and seeds.

The impressive degree of coordination between the decrease in the level of *VvNCED1* transcript and the increase in the level of *VvA8H-CYP707A4* transcript in response to dormancy release stimuli, which we recorded in grapevine buds as described above, may serve as a hint for the potential existence of a shared regulatory entity that controls both, and thereby serves as a critical upstream switch, itself being regulated by dormancy release stimuli. Coordinated regulation of *NCED* and *CYP707A* gene expression in response to unfavorable light or temperature conditions has been recorded in seeds of several species (Argyris *et al*., 2011; Gubler *et al*., 2008; Leymarie *et al*., 2009; Seo *et al*., 2006; Toh *et al*., 2008). Interestingly, in Arabidopsis, key genes for ABA metabolism and signaling are highly sensitive to slow seasonal changes in soil temperature, and regulation of their expression results in continual and dramatic adjustments to the dormancy depth within the soil seed bank. In relation to our focus, low soil temperature (which is linked to deep seed dormancy) upregulated expression of *AtNCED6*, whereas the transition to high soil temperature (which induces dormancy release) was accompanied by increased expression of *AtCYP707A2* (Footitt *et al*., 2011). A striking mechanistic understanding of the integrated regulation of ABA synthesis and catabolism genes was recently achieved in rice seeds. The molecular switch in this case appears to comprise two interacting bHLH TFs (basic/helix-loop-helix transcription factors), OsSD6 (Seed Dormancy 6) and OsICE2 (inducer of C-repeat binding factors expression 2), which antagonistically control seed dormancy (Xu *et al*., 2022). Whereas OsSD6, the causal gene within an identified seed dormancy QTL, negatively regulates rice seed dormancy, OsICE2 plays a positive role in regulating seed dormancy. Excitingly, OsSD6 and OsICE2, which physically interact, regulate dormancy by oppositely regulating *OsNCED2* and *OsABA8OX3* expression. While regulation of *OsABA8OX3* expression by OsSD6-OsICE2 is direct, the regulation of *OsNCED2* expression is indirect, being exerted via regulation of a third bHLH TF, OsbHLH048, a direct repressor of *NCED* expression. In addition, it was shown that this regulatory “duo” operates in a temperature-dependent manner. Soil temperatures above 25 °C, which are suitable for germination, elevate the expression of *OsSD6*, but sharply reduce expression of *OsICE2*. In contrast, low temperatures, which inhibit germination, enhance *OsICE2* expression. A model was proposed in which (1) increased *OsSD6* expression and decreased *OsICE2* expression results in upregulation of *OsABA8OX3* and down regulation of *OsNCED2* (via upregulation of *OsbHLH048*), whereas (2) increases in, or high levels of *OsICE2* expression result in the opposite scenario, leading to increases in ABA content and induction of dormancy (Chen *et al*., 2023; Xu *et al*., 2022).

In light of the above, an assumption was raised that conservation of this newly exposed SD6-ICE2 regulatory module might exist and might play a role in the regulation of bud dormancy across woody perennials. Motivated by this assumption, we identified the potential homologs of OsSD6, OsICE2 and OsbHLH48 that are expressed in grapevine woody buds, and followed the modulation in their expression in response to dormancy release stimuli.

## MATERIALS AND METHODS

### Plant material

Mature buds were collected from grapevines (*Vitis vinifera* cv. Early sweet) in a commercial vineyard, at Gilgal located in the Jordan Valley, as previously described (Zheng *et al*., 2018a). Vines were pruned to three-node spurs, and the detached canes, each carrying nine buds (in positions 4-12), were cut into single-node cuttings and randomly mixed.

### Natural dormancy curve

Canes were sampled at several time points from November 2021 to January 2022, and nine groups of 10 single-node cuttings were placed in open vases containing tap water under the forcing condition (22°C with a 14 h/10 h light/dark regime). Bud break was monitored at 7, 10, 14, 17, 21, 24, and 28 d, and the bud break percentages at 21 d were used to prepare a seasonal dormancy curve, as described by Zheng *et al*. (2015). For gene expression, three groups of one hundred buds were randomly selected at each sampling point from the pool of single node cuttings, at the day of arrival from the vineyard. These bud pools were frozen in liquid nitrogen and stored at -80°C.

### Chemical and physical treatments

#### Analysis of the effect of HC on gene expression using an Open Vases (OV) experimental system

To test the effect of HC [3% (v/v) Dormex® (SKW, Trostberg, Germany)], single-node cuttings were treated as described in Zheng *et al*. (2015). Cuttings treated with 0.02% Triton X-100 served as controls. The treated cuttings were placed in open vases, under the forcing condition described above, and buds were sampled at 24, 48, and 96 h following treatment, frozen and stored at -80°C.

#### Analyses performed using a Sealed Vases (SV) experimental system

To validate our previous RNA-seq data, an enclosed environment was set up to test the effect of HC and hypoxia treatments on bud dormancy release. Treatments were carried out in sealed 2 L jars (three jars per treatment) as previously described (Ophir *et al*., 2009, Shi *et al*., 2018). In short, canes were sampled on Dec 17^th^, 2017, from the vineyard. Seven groups of 10 single-node cuttings per jar were immersed in 150 mL tap water. Cuttings were sealed for 24 or 48 h under the forcing condition (22°C under a 14/10 h light/dark), and then transferred to open vases. Buds were sampled from sealed jars separately at 24 or 48 h, frozen, and stored at -80°C.

To test the effect of HC in the enclosed system, single-node cuttings were sprayed with 3% Dormex®, and placed in jars as described above. For the relevant control treatment, cuttings were sprayed with 0.02% (v/v) Triton X-100 and incubated for 48 h in the sealed jars in the presence of a perforated tube containing vermiculite saturated with 7.43 g/100 mL KMnO_4_ solution (Merck, Darmstadt, Germany) as described by Ophir *et al*. (2009) and Shi *et al*. (2018). Buds were sampled as previously described.

Hypoxia treatments were set up in jars flushed with N_2_ (g) to reduce the O_2_ (g) level to 1%. The concentration of oxygen within the jar was determined using an OXYBABY 6.0 gas analyzer (WITT-Gasetechnik GmbH & Co. KG, Witten, Germany). Cuttings were sealed for 24 or 48 h under the forcing condition. This treatment was compared to the OV control. Other details are as previously described (Shi *et al*., 2020).

#### RNA extraction from buds and qRT-PCR analyses

Total RNA extraction and qRT-PCR were carried out as previously described (Acheampong *et al*., 2015; Acheampong *et al*., 2017; Zheng *et al*., 2015). The sequences of the gene-specific primers are provided in Table S1.

#### RNA-seq Data Availability

The Illumina RNA-seq raw data that were used to profile expression within the woody bud at 24 and 48 h after exposure to HC or hypoxia condition in the SV system (see *Shi et al.*, 2020) are available at the NCBI sequence-reads archive (SRA) under BioProject accession: PRJNA596362. The BioSample descriptions can be browsed using accession numbers SAMN13619444-57 and the raw data are accessible for downloading using accession numbers SRR10724800-13. All other relevant data can be found in Shi *et al*., 2020.

#### Phylogenetic Analysis

Phylogenetic analysis of bHLH proteins was performed using the full-length protein sequences. The protein sequences of 94 bHLH members in grape were reported by Wang *et al*. (2018). The sequences of AtICE1 (At3g26744), AtICE2 (At1g12860), AtbHLH57 (At4g01460), AtSPT (At4g36930), AtPIL5 (At2g20180), and AtZOU (At1g49770) in *Arabidopsis thaliana* were originally reported by Toledo-Ortiz *et al*. (2003). The sequences of OsSD6 (LOC_Os06g06900), OsICE1 (LOC_Os11g32100), OsICE2 (LOC_Os01g70310), and OsbHLH048 (LOC_Os02g52190) in *Oryza sativa* were obtained from Xu *et al*. (2022). All bHLH proteins were initially aligned with ClustalW, and the MEGA X software was used to conduct a phylogenetic analysis based on the neighbor-joining method on 1000 bootstrap replications with the default parameters (Kumar *et al*., 2018).

## RESULTS

To identify the potential homologs of OsSD6, OsICE2 and OsbHLH48 that are expressed in grapevine woody buds, we first constructed a phylogenetic tree comprising the relevant genes from rice {OsSD6 (LOC_Os06g06900), OsICE2 (LOC_Os01g70310), OsICE1 (LOC_Os11g32100), OsbHLH048 (LOC_Os02g52190)}, their homologs from arabidopsis {AtICE1 (AT3G26744), AtICE2 (AT1G12860) and AtbHLH57 (AT4G01460)}, and the 94 members of the bHLH family in grape (Wang *et al*., 2018). This analysis revealed that (1) VvbHLH85 and VvbHLH29 (VvSD6) are orthologous to OsSD6, (2) VvbHLH074 (VvICE2), VvbHLH060 (VvICE1-1) and VvbHLH06 (VvICE1-2) are orthologous to OsICE1, OsICE2, AtICE1, and AtICE2, (3) VvbHLH067, VvbHLH064, VvbHLH035, VvbHLH080 and VvbHLH08 are orthologous to OsbHLH048 and AtbHLH057 (Fig. 1). Since only *VvbHLH08*, *VvbHLH80*, *VvbHLH64* were expressed in the grape buds (Shi *et al*., 2020) and presented highly similar expression profile (Fig. S1), VvbHLH64 was selected as the representative homolog of OsbHLH048 for further analyses.

**Figure 1.**
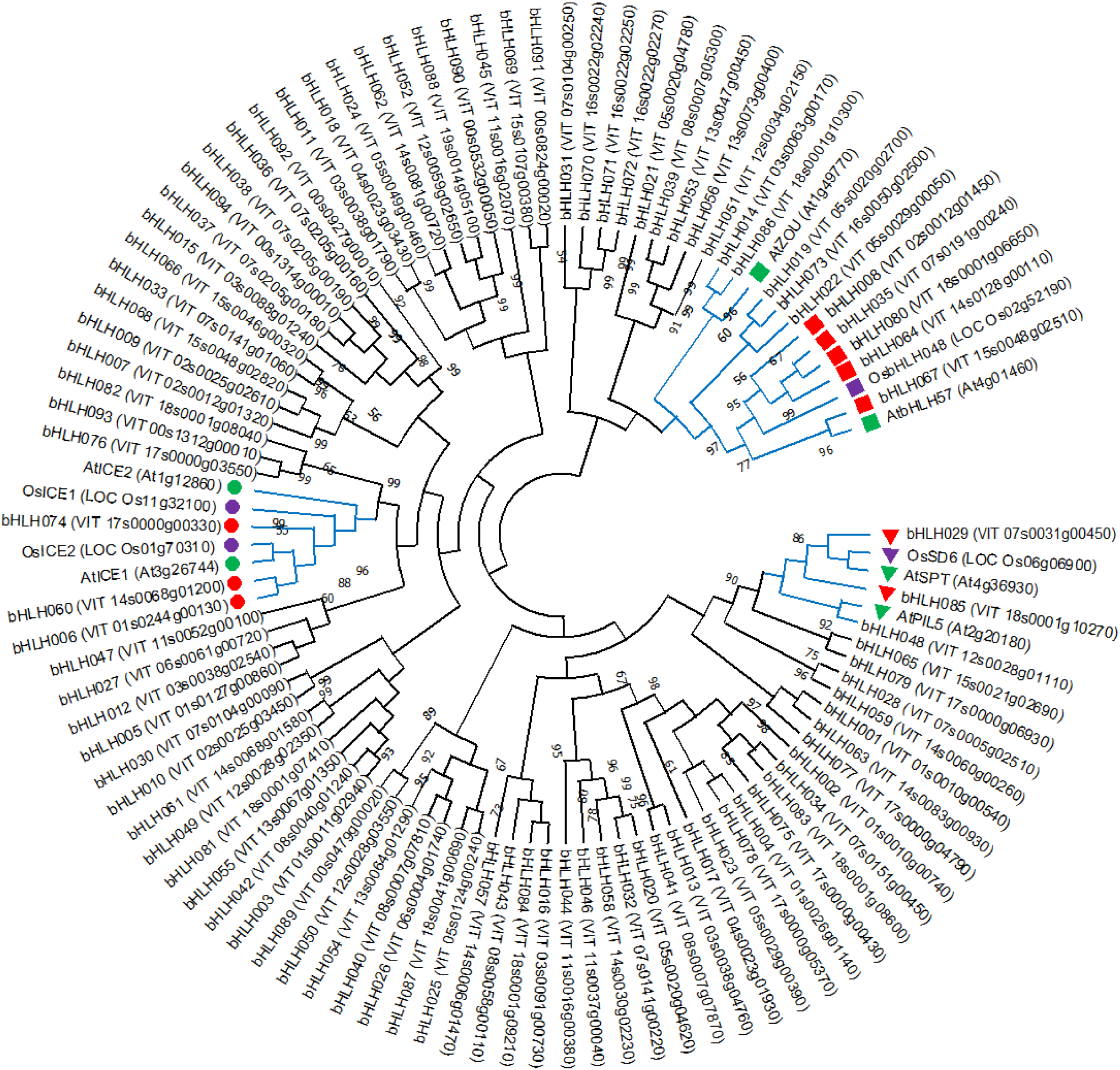
Neighbour-joining tree based on the amino acid sequence alignment of OsSD6, OsICE1, OsICE2 and OsbHLH048 from *Oryza sativa*, AtSPT, AtICE1, AtICE2, AtbHLH057, AtZUO and AtPIL5 from *Arabidopsis thaliana* and the members of the bHLH family in *Vitis vinifera*. Full-length protein sequences of 94 bHLH members of *Vitis vinifera*, and selected bHLH proteins from *Oryza sativa* (OsSD6, OsICE1, OsICE2 and OsbHLH048) and *Arabidopsis* (AtSPT, AtICE1, AtICE2, AtbHLH57, AtZOU and AtPIL5) were aligned using the CLUSTAL-W algorithm of MEGA-X software (version 10.0, Kumar *et al*., 2018). The aligned sequences were then used to reconstruct phylogenetic tree using the neighbor-joining method (MEGA X software). Bootstrap values as a percentage of 1000 replicates were calculated for each node. Bootstrap values less than 50 are not shown. Square symbols indicate the OsbHLH048 subfamily, inverted triangle symbols indicate the SD6 subfamily, and circle symbols indicate the ICE subfamily. The red, purple and green markers represent the relevant bHLH members from *Vitis vinifera*, and *Oryza sativa*, and *Arabidopsis thaliana*, respectively.

To assess the potential roles of these candidates in dormancy release of grapevine buds, we initially retrieved their expression profiles from a transcriptomic database that we previously generated (Shi *et al*., 2020). These data report the effect of HC on buds carried on woody single-node cuttings that were placed in sealed jars for 48 h under forcing condition (Sealed Vase system, SV). According to these RNAseq data (Fig. 2A-F), the orthologues of *OsSD6* (*VvbHLH85* and *VvSD6*) are both upregulated in response to HC. The ICE orthologues differ in their response. Expression of the orthologue of *OsICE2* (*VvICE2*), is down regulated, the expression of one orthologue of *OsICE1* (*VvICE1-2*) is upregulated, while that of the other (*VvICE1-1*) is not affected by HC.

**Figure 2.**
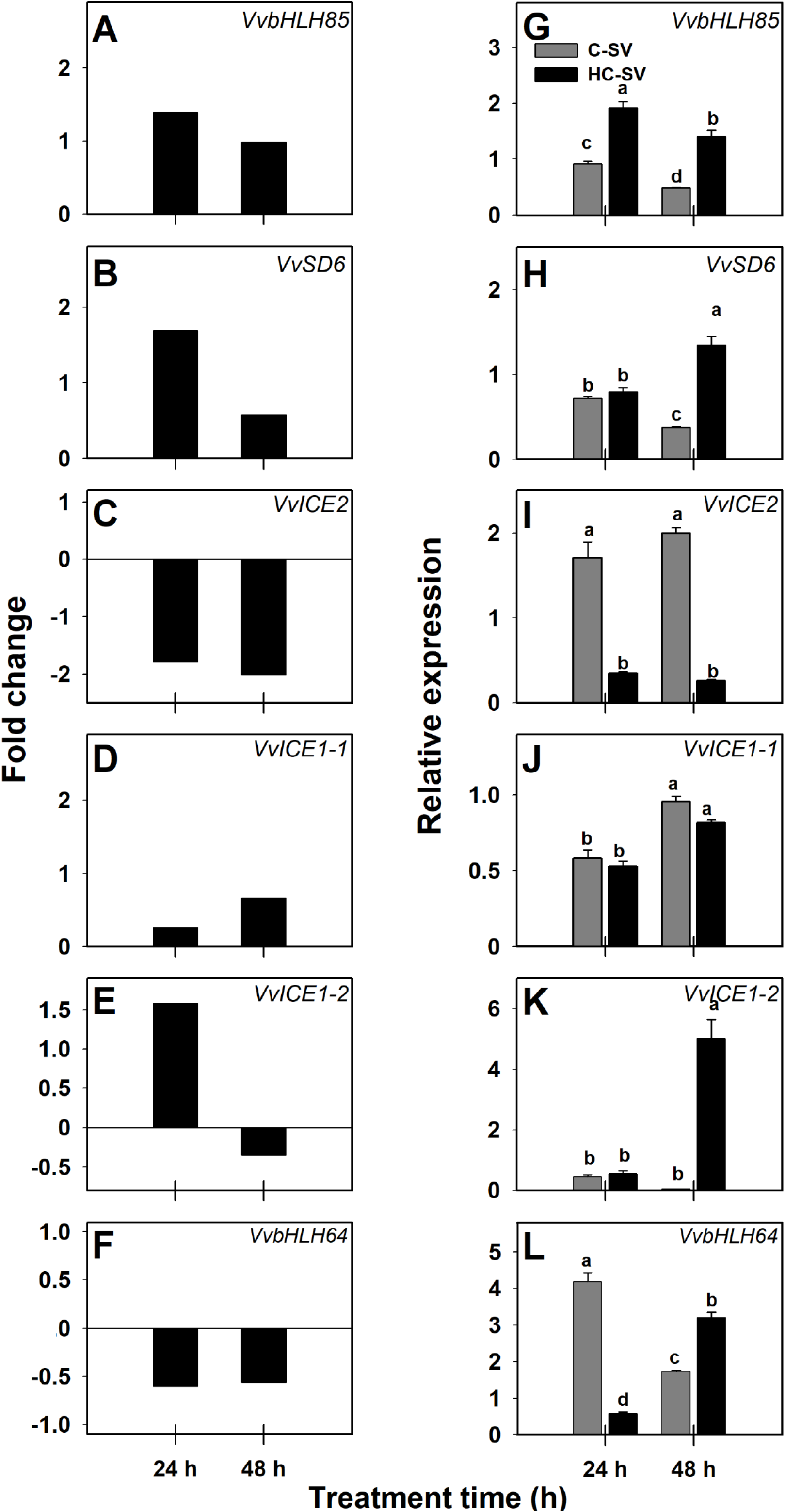
The effect of HC on the expression of six *VvbHLH* genes that are orthologous to *OsSD6*, *OsICE2*, and *OsbHLH048* in grapevine buds placed under SV conditions. (A-F) RNA-seq based expression profiles are presented for *VvbHLH85* (A), *VvSD6 (VvbHLH29*, B), *VvICE2* (*VvbHLH74*, C), *VvICE1-1* (*VvbHLH60*, D), *VvICE1-2* (*VvbHLH06*, E), and *VvbHLH64* (F). Single-node cuttings were sprayed with 3% Dormex®, and placed in 2 L jars (three jars per treatment, seven groups of 10 cuttings per jar, immersed in 150 mL water). The jars were sealed for 24 and 48 h under the forcing condition (22°C under a 14/10 h light/dark). For the relevant control treatment, cuttings were sprayed with 0.02% (v/v) Triton X-100 and incubated for 24 or 48 h as described above, in the presence of a perforated tube containing vermiculite saturated with 7.43 g 100 ml^-1^ KMnO_4_ solution. Total RNA was extracted from HC-treated buds and KMnO_4_-control buds sampled at 24 and 48 h after treatment, and was used for RNA-seq analysis (reported in Shi *et al*., 2020). The raw RNA-seq data used to profile expression is available at the NCBI sequence-reads archive (SRA) under BioProject accession: PRJNA596362. (G-L) Canes were collected on Dec 17^th^, 2017. Single-node cuttings were treated with 3% HC and 0.02% (v/v) Triton X-100 with KMnO_4_ solution in sealed jars as described above. 15 out of 70 buds were used for RNA extraction. Relative transcript levels were determined for *VvbHLH85* (G), *VvSD6* (H), *VvICE2* (I), *VvICE1-1* (J), *VvICE1-2* (K), and *VvbHLH64* (L), using qRT-PCR normalized against *VvActin*. The qRT-PCR values are the means of two biological replications, each with three technical repetitions, ± SE. Statistical tests indicate differences between treatments in each time point. Data points with different letters indicate values that are significantly different (*P* < 0.05) according to Tukey’s HSD test. For additional details, see Experimental Procedures and Table S1.

To verify these results, qRT-PCR analyses were carried out, using identical treatments and experimental system (HC; SV system), but with plant material collected in a different year (see M&M). The qRT-PCR results fully confirmed the direction of regulation reported above, suggesting (1) significant up-regulation of the grapevine orthologues of *OsSD6* (*VvbHLH85* and *VvSD6*) as well as *VvICE1-2* at 48 h after HC application (Fig. 2G, H, K), (2) significant down-regulation of *VvICE2* (Fig. 2I), which like *VvbHLH85* is evident already at 24 h after HC application, (3) no effect of HC on *VvICE1-1* (Fig. 2J). A mixed direction of regulation was recorded for *VvbHLH64*, in which significant down-regulation at 24 h was followed by upregulation at 48 h (Fig. 2L), which does not agree with the RNAseq data (Fig. 2F).

We then studied whether such solid and opposite effects of HC on the expression of *VvSD6* and *VvICE2* orthologues could be detected using our Open Vase experimental system (OV), in which the single-node cuttings are placed in open jars under identical forcing conditions (Shi *et al*., 2018), and expression is monitored for 4 d. In this analysis, we compared the expression profiles in buds sampled at three different dates (November 27, December 4. and December 11) over the natural dormancy cycle (Fig. S2). According to the results, both homologs of *OsSD6* are significantly upregulated at 48 h at all three sampling dates (Fig. 3A-F). Whereas *VvbHLH85* expression is significantly upregulated at 24 h as well, for all three sampling dates, upregulation of *VvSD6* at 24 h is significant only on December 4. At 96 h, there is no significant difference in transcript levels between HC and control samples for both genes at all three tested sampling dates. Similar to *VvbHLH85* and *VvSD6*, the expression of *VvICE1-2* was significantly upregulated at 24 h (November 27 and December 11, Fig. 3M, O) or 48 h (December 4, Fig. 3N), whereas expression of *VvICE1-1* is not significantly regulated by HC (Fig. 3J-L). Similar to its behavior in the SV system (Fig. 2C, I), the level of the *VvICE2* transcript is considerably down regulated at both 24 and 48 h for all three sampling dates (Fig. 3G-I). A trend of down-regulation appears at 96 h, being significant only on December 11 (Fig. 3I). As for *VvbHLH64*, the data do not support a clear pattern of regulation of expression by HC (Fig. 3P-Q), significant upregulation being recorded in buds sampled on December 11 representing 48 h after HC application (Fig. 3R). Based on the above, it appears that, similar to the situation during rice germination, HC application that induce bud dormancy release, result in coordinated up-regulation of *OsSD6* homologues and down-regulation of *OsICE2* homologues.

**Figure 3.**
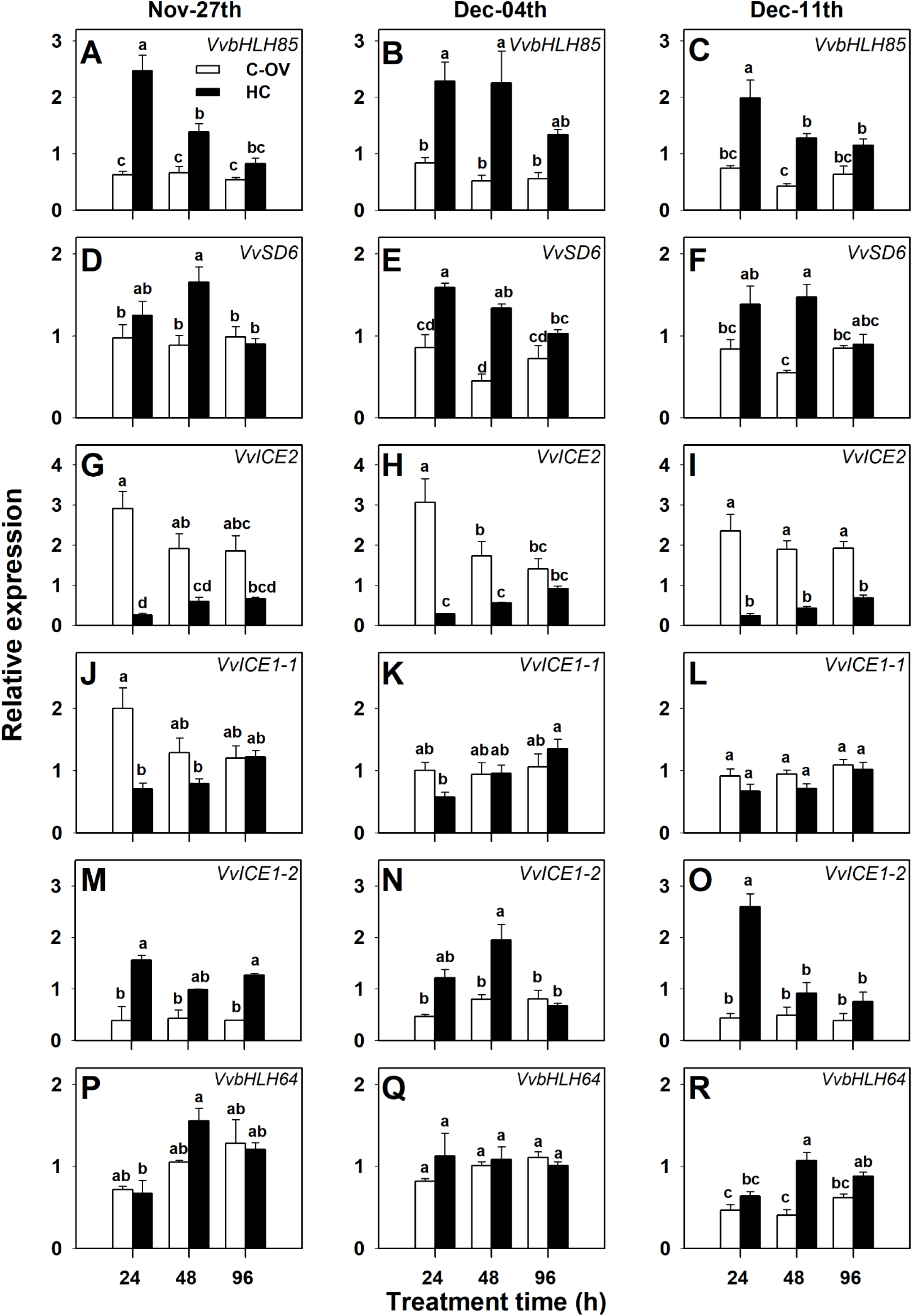
The effect of HC on the expression of *VvbHLH85*, *VvSD6, VvICE2, VvICE1-1, VvICE1-2* and *VvbHLH64* in grapevine buds placed under OV conditions at three time points during the dormancy cycle. (A-R) Canes were collected on Nov 27^th^, Dec 04^th^ and Dec 11^th^ in 2021. Seven groups of 10 single-node cuttings were spray-treated with HC, as described in Fig. 2, with a similar number of cuttings treated with 0.02% Triton X-100 serving as controls. The treated cuttings were immersed in open vases containing 150 ml tap water and placed under the forcing condition (22°C with a 14 h/10 h light/dark regime). Buds were sampled at 24, 48, and 96 h following treatment. Total RNA was extracted and the relative transcript levels determined for *VvbHLH85* (A-C), *VvSD6* (D-F), *VvICE2* (G-I), *VvICE1-1* (J-L), *VvICE1-2* (M-O), and *VvbHLH64* (P-R), using qRT-PCR normalized against *VvActin*. The qRT-PCR values are the means of two biological replications, each with three technical repetitions, ± SE. For additional details, see Experimental Procedures and Figure 2.

Comparison of the response of the buds to HC to its response to a different stimulus of bud dormancy release is critical to provide support for the assumption that regulation of expression of the *SD6*–*ICE* duo is correlated to bud dormancy release. Accordingly, we analyzed the influence of hypoxia on the expression profile of all the genes reported above. Both the transcriptomic data (originated from a data set published by Shi *et al*., 2020) and the qRT-PCR results (Fig. 4), collected using buds sampled over two different years, are presented. In agreement with the responses to HC, the data reveal remarkable upregulation of both orthologues of *OsSD6* (*VvbHLH85* and *VvSD6*) and of *VvICE1-2* at 24 and 48 h after exposure to hypoxia (Fig. 4A, B, E, G, H, K). Similarly, they display down-regulation of *VvICE2* at 48 h (Fig. 4C, I). Of special interest are the profiles of *VvICE1-1* and *VvbHLH64*. The expression levels of these genes, which was not regulated by HC, were considerably and significantly upregulated and downregulated by hypoxia, respectively (Fig. 4D, F, J and L).

**Figure 4.**
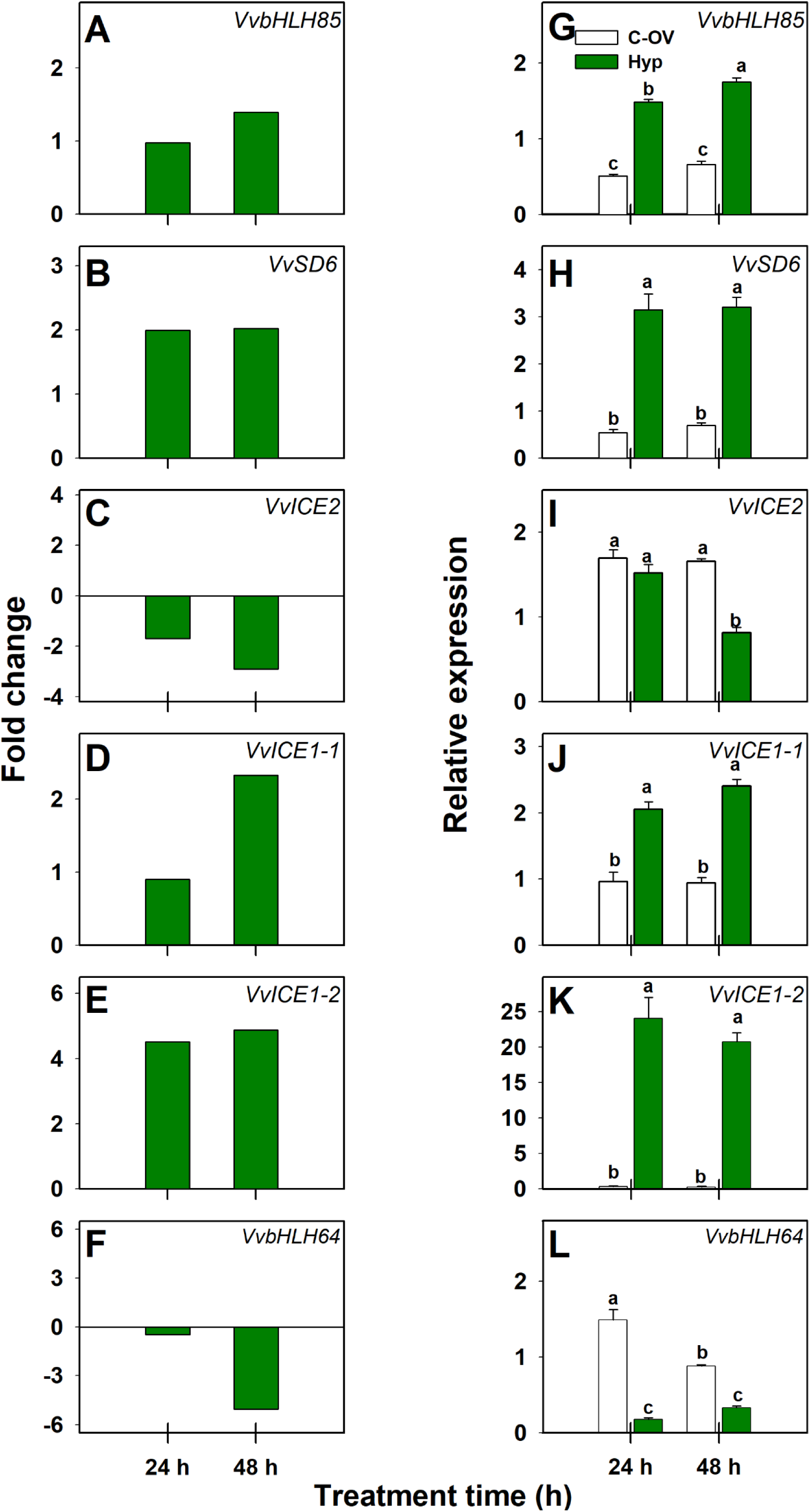
The effect of hypoxia on the expression of *VvbHLH85*, *VvSD6, VvICE2, VvICE1-1, VvICE1-2,* and *VvbHLH64* in grapevine buds. (A-F) Hypoxia treatment (1% O_2_) was set up in sealed jars flushed with N_2_ (see Fig. 2 for experimental SV setting). Buds were sampled from the sealed jars at 24 and 48 h. Cuttings in open vases were sampled in parallel to serve as controls. Total RNA was extracted from hypoxia-treated and control buds sampled at 24 and 48 h after treatment, and was used for RNA-seq analysis (see details in Fig. 2). RNA-seq based expression profiles of hypoxia treatment are presented for *VvbHLH85* (A), *VvSD6 (*B), *VvICE2* (C), *VvICE1-1* (D), *VvICE1-2* (E), and *VvbHLH64* (F). (G-L) Canes were collected on Dec 17^th^, 2017. Hypoxia treatments and relevant controls were set up as described above. 15 out of 70 buds were used for RNA extraction. Relative transcript levels of *VvbHLH85* (G), *VvSD6* (H), *VvICE2* (I), *VvICE1-1* (J), *VvICE1-2* (K), and *VvbHLH64* (L) were determined for hypoxia and untreated buds, using qRT-PCR normalized against *VvActin*. For additional details, see Experimental Procedures and Figure 2.

To chart the direction of regulation of the genes under study over the natural dormancy cycle, we profiled their transcript levels in buds sampled at various time points, and described these in terms of dormancy depth. Based on the data, the profiles can be divided into two principal groups (Fig. 5). In the case of *VvSD6* and *VvICE1-2*, the transcript levels were low during dormancy induction, significantly increasing at maximal dormancy and remaining high throughout dormancy release (Fig. 5A, B, E). On the other hand, expression of *VvICE2* and *VvbHLH64*, similarly increased during dormancy induction, reaching a maximum during the period of deepest dormancy, and presenting a significant decrease accompanying dormancy release (Fig. 5C, F). The expression level of *VvICE1-1* presented no change during dormancy induction, and a slight, yet significant down-regulation during dormancy release (Fig. 5D).

**Figure 5.**
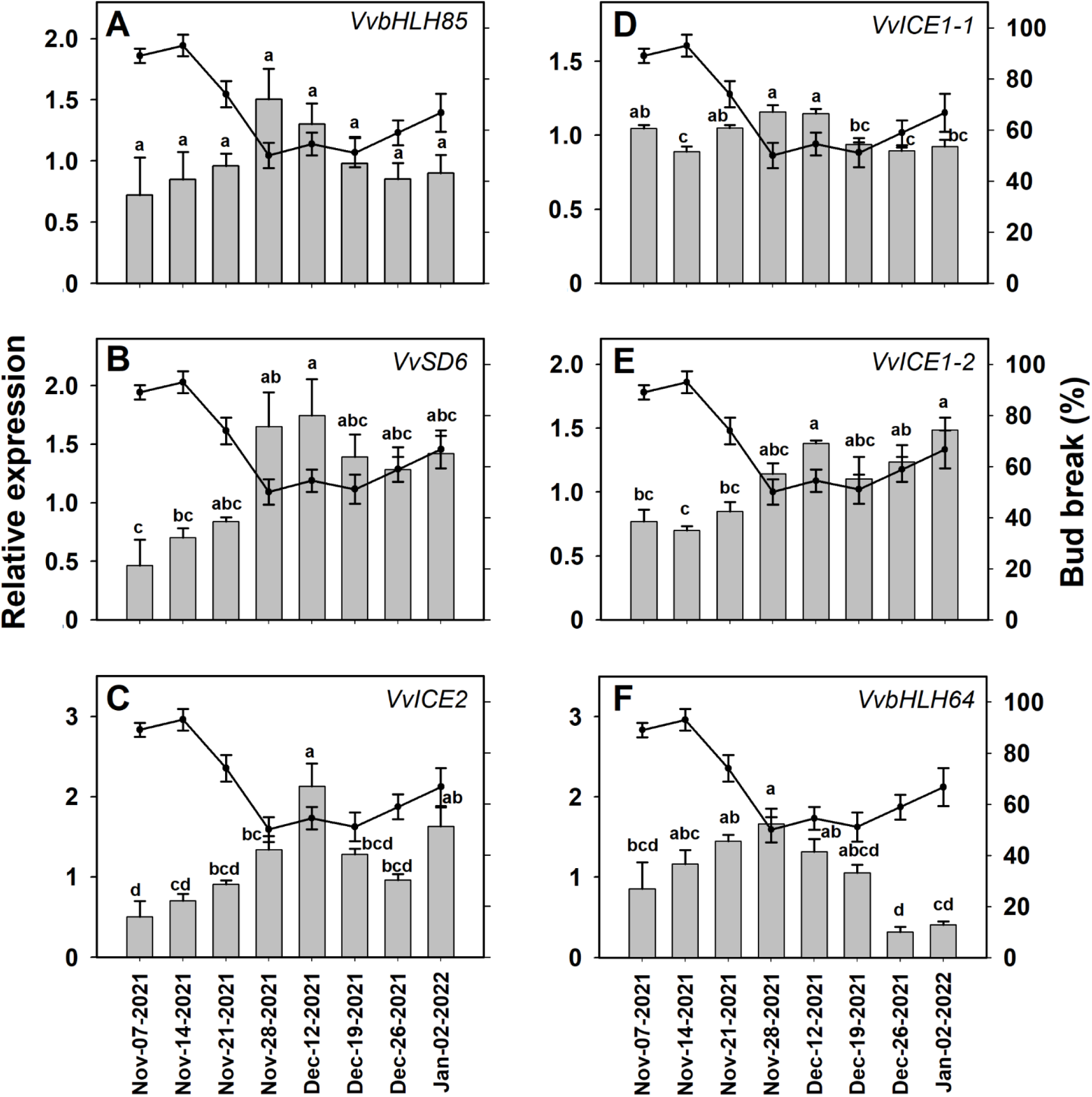
Profiling transcript levels of *VvbHLH85*, *VvSD6, VvICE2, VvICE1-1, VvICE1- 2,* and *VvbHLH64* in grapevine buds throughout the dormancy cycle. Canes were sampled weekly at eight time points from November 2021 to January 2022. Single-node cuttings were randomly mixed. Nine groups of 10 single-node cuttings were placed in open vases containing tap water under the forcing condition (22°C with a 14 h/10 h light/dark regime), and used for bud break monitoring. The bud-break percentages at 21 days are shown (as a line) in each panel. Values are averages of nine groups of 10 buds each, ± SE. Total RNA was extracted from 15 buds sampled weekly, upon arrival from the vineyard, and frozen. Relative transcript levels were determined for *VvbHLH85* (A), *VvSD6* (B), *VvICE2* (C), *VvICE1-1* (D), *VvICE1-2* (E) and *VvbHLH64* (F), using qRT-PCR normalized against *VvActin*. For additional details, see Experimental Procedures and Figure 2.

The bHLH-type TFs often bind to E-box (CANNTG) or G-box (CACGTG) motifs located in the promoters of target genes (Chaudhary and Skinner, 1999; Oh *et al*., 2007). In agreement, OsSD6 and OsICE2 from rice function antagonistically in controlling seed dormancy by interacting with G-boxes or E-boxes in the promoter of *OsABA8OX3* and *OsNCED2* (Xu *et al*., 2022). We therefore examined whether such motifs exist in the promoters of the ABA biosynthesis gene *VvNCED1* and the ABA catabolism gene *VvA8H- CYP707A4*. We identified two G-boxes in the promoters of *VvNCED1* (-873 and -862 bp upstream of the first codon) and of *VvA8H-CYP707A4* (-409 and -292 bp upstream of the first codon, Fig. 6).

**Figure 6.**
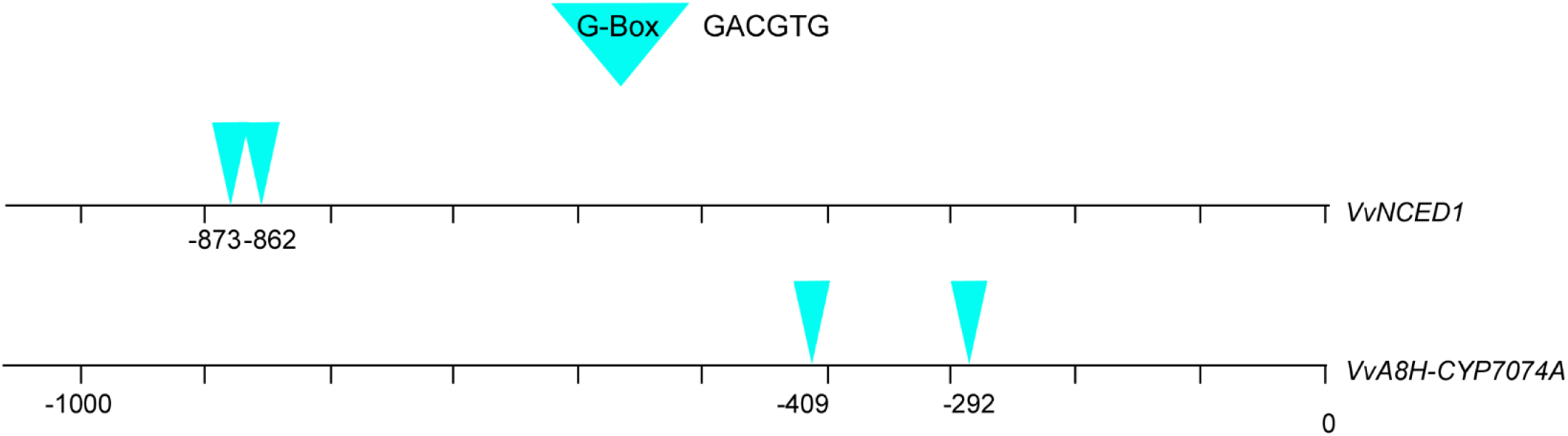
**Analysis of the promoter sequences of the *VvNCED1* and *VvA8H-CYP707A4*** genes for bHLH-type transcription factor binding sites. Nucleotide sequence between positions − 2000 to 0 bp upstream from the transcription start site (ATG) of *VvNCED1* (Vitvi05g00963) and *VvA8H-CYP707A4* (Vitvi03g00508) genes were downloaded from the *Vitis vinifera* (PN40024.v4) genomic database (*Vitis Vinifera*- Ensembl Genomes 51, https://plants.ensembl.org/Vitis_vinifera/Info/Index) according to the gene localization. PlantCARE (http://bioinformatics.psb.ugent.be/webtools/plantcare/html/) was used to analyze the G-box (GACGTG) and/or E-box (CANNTG) motifs in the promoter region of *VvNCED1* and *VvA8H-CYP707A4* genes (Lescot *et al*., 2002). Two G-boxes are located at -873 bp and -862 bp upstream of ATG of *VvNCED1*. Another two G-boxes are located at -409 bp and -292 bp of the promoter region of *VvA8H-CYP707A4*. The triangles and the numbers indicates the location of G-box motifs.

## DISCUSSION

We have previously shown that treatment with exogenous ABA delays dormancy release in grapevine buds (Zheng *et al*., 2015). Observation of this inhibitory effect suggested that ABA might be an important component of the regulatory network that normally operates during the dormancy cycle in grapevine buds. Hence, we proposed that both natural and artificial stimuli of bud dormancy release might exert their effects by decreasing endogenous ABA levels within the buds. We also proposed that such reduction in ABA levels is achieved, at least partially, via upregulation of *VvA8H-CYP707A4,* the highest bud-expressed paralog of the gene family encoding ABA degrading enzymes, ABA 8’-hydroxylase (ABA8’OH) (Fasoli *et al*., 2012; Zheng *et al*., 2015). In agreement with this proposition, we found that (1) several artificial stimuli of dormancy-release (HC, HS, AZ), that trigger respiratory stress, induce *VvA8H- CYP707A4* expression; (2) the *VvA8H-CYP707A4* transcript level is also modulated during the natural dormancy cycle, and markedly increased during the transition from the deep dormancy phase to the dormancy release phase (Zheng *et al*., 2015).

In light of the above, and based on the recent discovery of the coordinated function of the SD6-ICE2 duo as direct regulators of *OsABA8OX3* transcription and germination of rice seeds, the objectives of the current study were to examine whether (1) expression of the *Vitis* homologs of rice SD6 and ICE2 is regulated by dormancy release stimuli, and (2) the direction of regulation is in agreement with the direction of regulation of *VvA8H-CYP707A4*.

Excitingly, we recorded significant upregulation of the two *VvSD6* homologs, and down-regulation of *VvICE2* in response to application of HC, which induces ABA8’OH gene expression and dormancy release in grape buds. The response is evident 24 h after treatment, and lasts longer in the case of *VvICE2* than for the *VvSD6* homologues (Fig. 2G-I, Table S2). Interestingly, similar changes were recorded in response to hypoxia (Fig. 4G-I, Table S2), which also strongly induces dormancy release, supporting a causal relationship between these TFs and bud dormancy release. The seasonal profile, which reflects a much wider window of time, may be divided to two periods roughly corresponding to a gradual entrance to dormancy followed by its release. The profiles of *VvSD6* (Fig. 5B, Table S2) and *VvICE2* (Fig. 5C, Table S2) presents upregulation and down regulation, respectively, from the stage of maximal dormancy. These findings further support the idea that the antagonistic expression profiles of *VvSD6* and *VvICE2* are correlated with ABA metabolism and bud dormancy release.

While the above perfectly follow the scenario described for the rice model, differences were evident in direction of regulation of the homolog of OsbHLH048 (VvbHLH64). In rice, OsSD6 activates *OsbHLH048* expression via direct binding to a G-box within its promoter. This results in inhibition of *OsNCED2*, and has a positive effect on germination. Oppositely, OsICE2 inhibits *OsbHLH048* expression via direct binding to an E-box within the same promoter, which results in enhancement of *OsNCED2* expression and has a negative effect on germination (Xu *et al*., 2022, Table S2). Since *OsbHLH048* serves as a repressor of *OsNCED2* transcription in rice, we expected an upregulation of its homolog (*VvbHLH64*), in response to dormancy release stimuli. However, in grapevine buds, it was either down-regulated, in response to hypoxia and during natural dormancy release, or not affected, in response to HC (Fig. 3P-R, Fig. 4L, Table S2). This finding raises the possibility that, in grape buds, *VvbHLH64* serves as an activator of *NCED* expression rather than a repressor.

While bHLH proteins from the three subfamilies discussed above are also involved in regulation of seed dormancy in Arabidopsis, the regulation of specific homologs is not necessarily in the same direction, the regulatory circuits appearing to be more complex in Arabidopsis and involving additional regulators. Interestingly, *AtbHLH57*, a homolog of the *OsbHLH048* that binds to *AtNCED6*/*9* promoter, serves as an activator of *AtNCED6*/*9* expression, rather than as a repressor as in rice (Liu *et al*., 2020; Xu *et al*., 2022). This direction of regulation agrees with the down-regulation of *VvbHLH64* during bud dormancy release in grape (Table S2).

While members of the *ICE* subfamily regulate *NCED* transcription indirectly in both rice and Arabidopsis, via regulating the transcription of *OsbHLH048*/*AtbHLH57* which binds to the *NCED* promotor, the cascade in Arabidopsis presents more layers resulting in action in the opposite direction. In short, seed-specific AtODR1 interacts with AtbHLH57 and inhibits AtbHLH57-modulated *AtNCED6/9* expression in the nucleus (Liu *et al*., 2020). Expression of AtABI3 (ABA INSENSITIVE3), which directly suppresses *AtODR1* expression (and thereby allows AtbHLH57-modulated At*NCED6/9* expression) is negatively controlled by a pair of bHLH proteins, AtICE1 and ZOU (which map to the subfamily of AtbHLH57, OsbHLH048, VvbHLH64, Fig. 1), that bind to the *AtABI3* promoter (MacGregor *et al*., 2019). Hence, AtICE1 is a negative regulator of *AtNCED6/9* expression and a positive regulator of arabidopsis seed germination, whereas rice OsICE2 is a positive regulator of *OsNCED2* and rice seed dormancy

The limited information available for AtICE2 suggests that it was derived from AtICE1 duplication, and acquired some unique features such as enhanced meristem freezing tolerance, high expression in axillary buds, presence of a SREATMSD conserved site which is considered a sugar responsive element (SRE), and presence of a RAP2.2 binding site, which is involved with hypoxia survival (Kurbidaeva *et al*., 2014).

Whereas the rice model involves only OsICE2 and the Arabidopsis model considers only AtICE1, which act oppositely in regulation of *NCED* expression, we recorded differential and opposite regulation of two members of the ICE subfamily, out of the three that are expressed in the bud (Table S2). One member, which we term *VvICE2*, is strongly down-regulated in response to HC, to hypoxia, and during natural dormancy release, and resembles the behavior of *OsICE2* (Fig. 2I, Fig. 3G-I, Fig. 4I, Fig. 5C). A second member, *VvICE1-1* was only regulated by hypoxia (Fig. 4J). The third member, *VvICE1-2*, is significantly up-regulated in response to all stimuli described above, in agreement with the Arabidopsis model (Fig. 2K, Fig. 3M-O, Fig. 4K, Fig. 5E). Notably, we could not detect expression of ABI3 and ABI4 homologs in the buds while expression of ABI5 homolog was not regulated by HC (Shi *et al*., 2020).

Based on integration of the literature with our data, and in light of the known function of bHLH TFs as homo-or heterodimers, it is tempting to speculate the existence of dynamic kinetics of dimer formation, where VvICE1-2 competes with VvICE2 on interaction with VvSD6. In this highly speculative scenario, VvICE2 interaction with VvSD6 will enhance *VvbHLH64* transcription, while VvICE1-2 – VvSD6 will synergistically inhibit *VvbHLH64* transcription.

It is worth noting at this stage that, as for *OsICE2* and *OsbHLH048*, *OsSD6* also has an Arabidopsis homolog, AtSPT (SPATULA). As in the case of OsSD6, a null mutation in AtSPT results in strong inhibition of germination of freshly matured seeds of the ecotype *Ler*, while AtSPT overexpression enhances germination. Rescue of the extreme *spt* dormancy phenotype was documented in response to inhibition of ABA synthesis, exogenous GA, decrease in endogenous ABA levels by *aba1* mutation, and removal of DELLA proteins (Vaistij *et al*., 2013). Despite this similarity, there is no record for AtSPT interaction with members of AtICE subfamily. Instead, AtSPT is repressed by AtSOM (SOMNUS), and is repressing the germination inhibitor AtMFT (MOTHER-OF-FT-AND-TFL1), two genes that are part of the phytochrome-controlled cascade that regulates seed germination (Vaistij *et al*., 2018)

The final conclusion in the context of the current study is the realization that bHLH TF genes that function as central regulators of ABA metabolism in Arabidopsis and rice seed probably serve as central regulators of ABA metabolism and dormancy release in buds of most if not all woody perennials. This powerful prediction, which should be confirmed in additional perennial systems, may serve as a handle to reveal factors upstream of the bHLH TFs that serve as primary inducers of bud dormancy release. A detailed analysis of the potential interactions between the genes and proteins addressed is warranted, and use of experimentally “friendly” bud systems, such as potato or poplar, may represent the best approach to provide a scaffold on which further understanding of upstream events may be assembled.

## SUPPLEMENTARY DATA

**Table S1.** Primers used in qRT-PCR analysis.

**Table S2.** Comparative summary of the regulation of *VvbHLH85*, *VvSD6, VvICE2, VvICE1-1, VvICE1-2*, and *VvbHLH64* by HC (in OV and SV) and hypoxia treatments, and during the natural dormancy cycle.

**Figure S1.** RNA-seq based expression profiles are presented for two orthologues of *VvbHLH64* in the dormant grape bud after HC and hypoxia-treatments in the SV experimental system.

**Figure S2.** Bud break analyses of HC-treated and control buds in the OV system at three sampling dates.

## AUTHOR CONTRIBUTIONS

Conceptualization: Zhaowan Shi, Sonika Pandey, Tamar Halaly-Basha, David W. Galbraith, Etti Or; Investigation: Zhaowan Shi, Sonika Pandey, Tamar Halaly-Basha; Writing-original draft: Zhaowan Shi, Etti Or; Writing-review & editing: David W. Galbraith, Etti Or.

